# Commensal rodent habitat expansion enhances arthropod disease vectors on a tropical volcanic island

**DOI:** 10.1101/2021.04.20.440633

**Authors:** De-Lun Wu, Han-Chun Shih, Jen-Kai Wang, Hwa-Jen Teng, Chi-Chien Kuo

## Abstract

Release from predators and competitors on volcanic islands can lead to increased body size and population density, as well as expanded habitat usage of introduced animals relative to their mainland counterparts. Such alterations might help the spread of diseases on islands when these exotic animals also carry pathogenic agents, but is rarely investigated. The commensal Asian house rat (*Rattus tanezumi*) is confined to human residential surroundings in Taiwan mainland but can be observed in the primary forests of the nearby Orchid Island, a tropical volcanic island. Orchid Island is also a hotspot of scrub typhus, a lethal febrile disease transmitted by larval trombiculid mites (chiggers) infective of the rickettsia *Orientia tsutsugamushi* (OT). We predicted an increase in chigger abundance when rodents, the primary host of chiggers, invade forests from human settlements as soils are largely devoid of in the latter habitat but is necessary for the survival of nymphal and adult mites. A trimonthly rodent survey in ten sites of three habitats (human resident, grassland, and forest) found only *R. tanezumi* and showed more *R. tanezumi* and chiggers in forest than in human residential sites. There was a positive association between rodent and chigger abundance, as well between rodent body weight and load of chiggers. Lastly, >95% of chiggers were *Leptotrombidium deliense* and their OT infection rates were similar among the habitats. Our study demonstrated potentially elevated risks of scrub typhus when the commensal rat is allowed to invade natural habitats on islands. In addition, while the success of invasive species can be ascribed to their parasites being left behind, island invaders might instead attain more parasites when the parasite requires only a single host (e.g. trombiculid mite), is a host generalist (*L. deliense*), and is transferred from unsuitable to suitable habitats (human settlement on mainland to forest on island).

## 1. Introduction

It is well acknowledged that biotas on insular oceanic islands are particularly vulnerable to invasion of exotic species compared to their continental counterparts. Most oceanic islands are depauperate in terrestrial biotas due to difficulty of species in colonizing isolated islands. This renders insular species less exposed to and thus evolving less defensive capabilities against predators and competitors (Paulay, 1994). Moreover, small land area leads to small population size and is therefore more susceptible to species extinction. Biological invasions can therefore have a devastating effect on island biodiversity (Reaser et al., 2007). For example, the introduction of a predatory snail to biologically control the invasive giant African snail has exterminated most endemic tree snail species in French Polynesia (Coote and Loeve, 2003). Bird species endemic to islands are more prone to extinction than continental ones, largely due to the detrimental effects of mammalian predators such as rats, cats, pigs etc. (Johnson and Stattersfield, 1990).

By contrast, little attention has been paid to whether oceanic islands can also facilitate spread of diseases when exotic species happen to carry pathogens or arthropod disease vectors. For example, rodents are common island invaders (Towns et al., 2006; Drake and Hunt, 2009) and also host many vector-borne and zoonotic diseases (Meerburg et al., 2009; Kosoy et al., 2015). At least three mechanisms can aid disease spread on islands (Fig. 1). First of all, ecological release from interspecific competition on species depauperate islands can allow habitat expansion of colonizers (Whittaker, 1998; Scott et al., 2003). For example, the flycatcher *Rhipidura fuliginosa* is confined to open habitats when sympatric with *R. spilodera* on the larger islands of New Hebridean, but also occurs in forests on the smaller islands where *R. spilodera* is absent (Diamond and Marshall, 1977). Likewise, the meadow vole occupies open fields on mainland but extend to woodlands on islands when the woodland-inhabited deer mouse occurs in the former but not the latter (Crowell, 1983). Disease-carrying invaders can therefore spread diseases following habitat expansion on oceanic islands, likely from human disturbed environments where most exotic species are introduced and adapt to (McKinney, 2002; Jesse et al., 2018), to more pristine habitats, like forests.

**Fig. 1.**
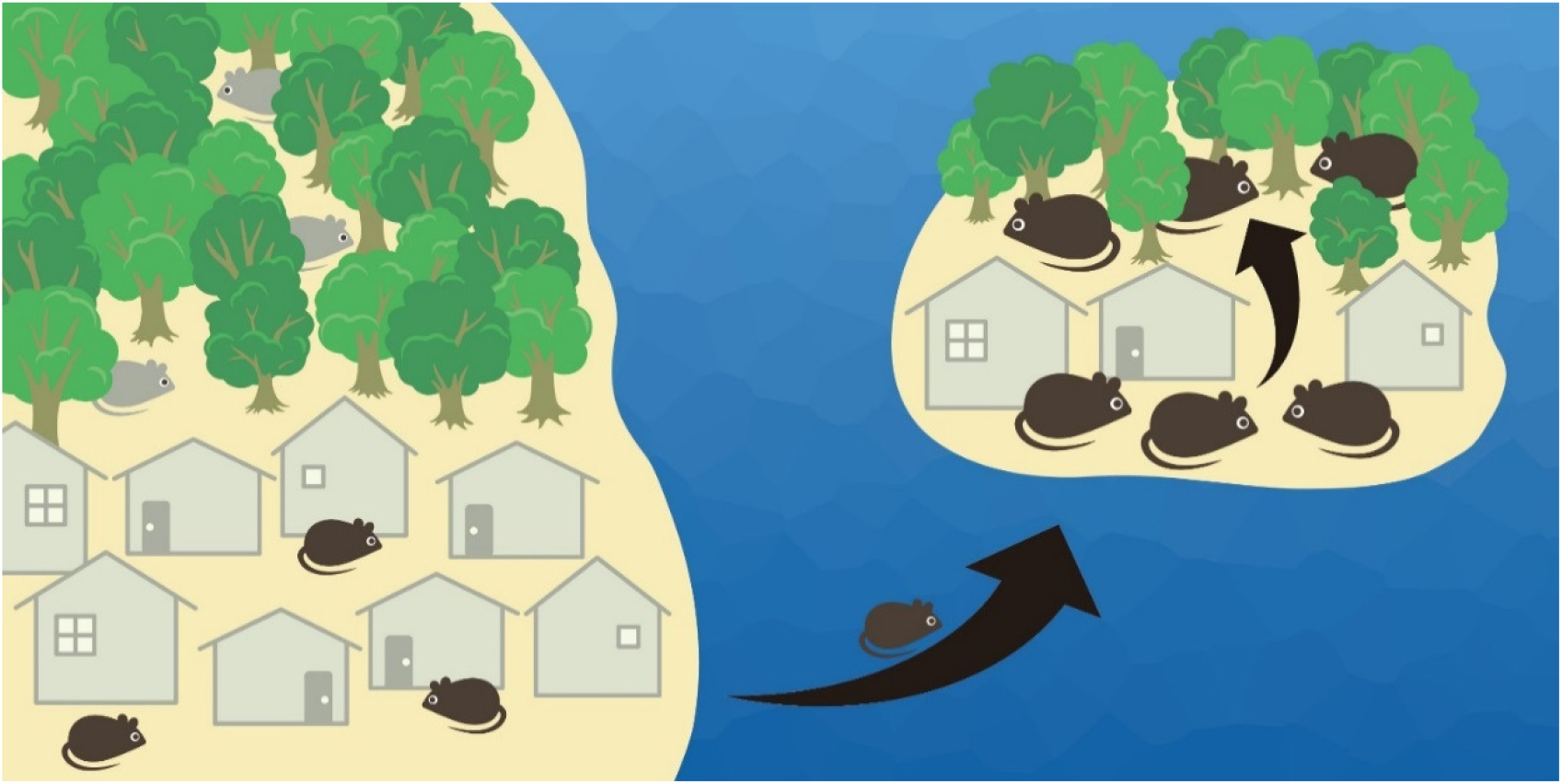
When colonizing ocean islands from mainland, rodents generally become larger and their population densities increase. Also, release from competitors on islands allow rodents to expand habitat usage; e.g. commensal rodent species (black color) is confined to human settlements on mainland when competitive species (gray color) occupies the forest, but is allowed to enter the forest when competitor is absent on islands.

Secondly, rodents often attain higher population density on islands than on mainland (Crowell, 1983; Adler and Levins, 1994; Adler, 1996). So called “island syndrome”, such systematic differences in demography is thought to be a direct effect of limited dispersal for insular populations (i.e. fence effect), as well as an indirect outcome of small land area that release island species from fewer predators and competitors (Adler and Levins, 1994). On the other hand, transmission of many pathogens are density dependent, with epidemiological models predicting that elevated density of susceptible host populations can increase contact rate and help sustain disease transmission (Swinton et al., 2002). For example, Sin Nombre virus was less detected after rodent population decline in the southwestern United States (Boone et al., 2002); although other correlation studies between host density and hantavirus prevalence don’t find a clear pattern, which is likely due to the limitations of antibody-based studies that can’t discern new from old infections (Forbes et al., 2018). Therefore, higher population density on oceanic islands could potentially facilitate disease transmission.

Lastly, rodent body size is usually larger on small islands than on mainland (Foster, 1964; Adler and Levins, 1994). Such gigantism could be the result of higher intra-specific competition following high population density on islands, thus favoring increased life span and body mass (Adler and Levins, 1994). This could also be due to the advantage of larger body size in securing food resources when small body size for escape from predation is not selected for on predator-lacked islands (Michaux et al., 2002). At the same time, it has been found that burden of some disease vectors (such as ticks) would increase with body mass of hosts (such as rodents) (Perkins et al., 2003; Kiffner et al., 2011; Mysterud et al., 2015). This is likely the results of larger surface areas for ectoparasites to attach for (Kuris et al., 1980) or larger hosts are more tolerant of parasite infestations (Hart et al. 1992; Olubayo et al. 1993). Consequently, increased body size of hosts on oceanic islands could presumably maintain a higher number of vectors and therefore help sustain vector-borne diseases.

Orchid Island, also known as the Lanyu Island, is a tropical volcanic island with an area of 46 km^2^ and 76 km off southeastern coast of Taiwan. The annual mean temperature was 22.8°C (monthly range: 18.6-26.2°C) and yearly rainfall totaled 2979 mm (average between 1991-2020, Taiwan Center Weather Bureau, https://www.cwb.gov.tw/V8/C/, assessed March 10, 2021). More than 80% of the Orchid Island is still covered with forests, mostly (ca. 80%) primary forests (Tsai et al., 2006). However, high abundance of the commensal Asian house rat (*Rattus tanezumi*) was observed in the primary forests and grassland of the Orchid Island (Shih, 2006). This is in contrast to the main Taiwan Island, where *R. tanezumi* is largely restricted to areas surrounding human settlements, as observed in other parts of Southeast Asia (Morand et al., 2015). Lowland forests and grassland are instead occupied by the native *Niviventer coninga* and *Rattus losea* (Lin, 1980).

Orchid Island is also a hotspot of scrub typhus. Scrub typhus is an acute and deadly infectious disease transmitted by larval trombiculid mites infective of the rickettsiae *Orientia tsutsugamushi* (OT). Previously confined to the western Pacific, southern Asia and northern Australia (Kelly et al., 2009), indigenous cases of this disease has lately been observed in South America (Weitzel et al., 2016; Claudine et al. 2017), Middle East (Izzard et al., 2010), and Africa (Thiga et al., 2015; Horton et al., 2016; Maina et al., 2016). At the same time, several countries have experienced a great increase in human incidences of scrub typhus (e.g. Roh et al., 2014; Wu et al., 2016; Sun et al., 2017). Life history of trombiculid mites includes egg, larva, nymph, and adult. Only the larval stage is parasitical, with rodents as the primary hosts (Harrison and Audy, 1951; Traub and Wisseman, 1974; Kawamura et al., 1995); mites in this stage are commonly called chiggers. The nymphal and adult stages are instead free living in the soils, predating on arthropods (Kawamura et al., 1995). Efficient transovarial and transstadial transmission of OT occurs in trombiculid mites, which are reservoirs of OT; animal hosts (e.g. rodents) play no role in transmitting OT (Kawamura et al., 1995). In eastern Taiwan where scrub typhus is already severe (Kuo et al., 2011), Orchid Island even tops among local districts in disease prevalence (Huang et al., 2009; Liu et al., 2014). In Orchid Island, antibody positivity rate for children at five and six years old was about 60% and 70-80%, respectively, but became 100% when children reached seven years old (Wu, 1993). Recently, an international traveler to the Orchid Island was died of scrub typhus (Wei et al., 2016).

Here, we investigated whether habitat expansion of the commensal Asian house rat will increase risks of scrub typhus in the extended habitats. Specifically, we hypothesized that chiggers will increase when the house rat expanded from human settlements to forests. This is due to that life cycle of trombiculid mites will be interrupted by paved grounds in the former habitat where soils necessary for nymphal and adult mites to survive are largely devoid of. In addition, although it has long been emphasized that scrub typhus can occur in a broad range of habitats other than scrub habitat, particularly forests (Traub and Wisseman, 1974), the relative suitability of forests in sustaining scrub typhus has never been quantified. Lastly, despite Orchid Island as a hotspot of scrub typhus, no systematic surveillance of chigger vectors on this island has ever been attempted; it was only briefly surveyed as part of a Taiwan-wide investigation on vectors of scrub typhus (Kuo et al., 2015). A good knowledge of chiggers, nevertheless, is helpful for prevention and control of scrub typhus on this island.

## 2. Material and methods

### 2.1. Study sites

We surveyed rodents and associated chiggers in 10 sites of three dominant habitat types (Lee et al., 2010) in the Orchid Island (Fig. 2). This included three human residential sites, three grassland sites, and four forest sites. Sites of the same habitat type were located in different regions of the island to control for potential regional difference in rodent and chigger abundance. Residential sites are characterized by paved-ground houses and roads surrounded with plant species adapted to human disturbance. Grassland is dominated by the silver grass (*Miscanthus sinensis*), while forests are composed of broadleaf species, mainly *Bischofia javanica, Ficus benjamina, Garcinia linii, Palaquium formosanum*, *Pometia pinnata* etc.

**Fig. 2.**
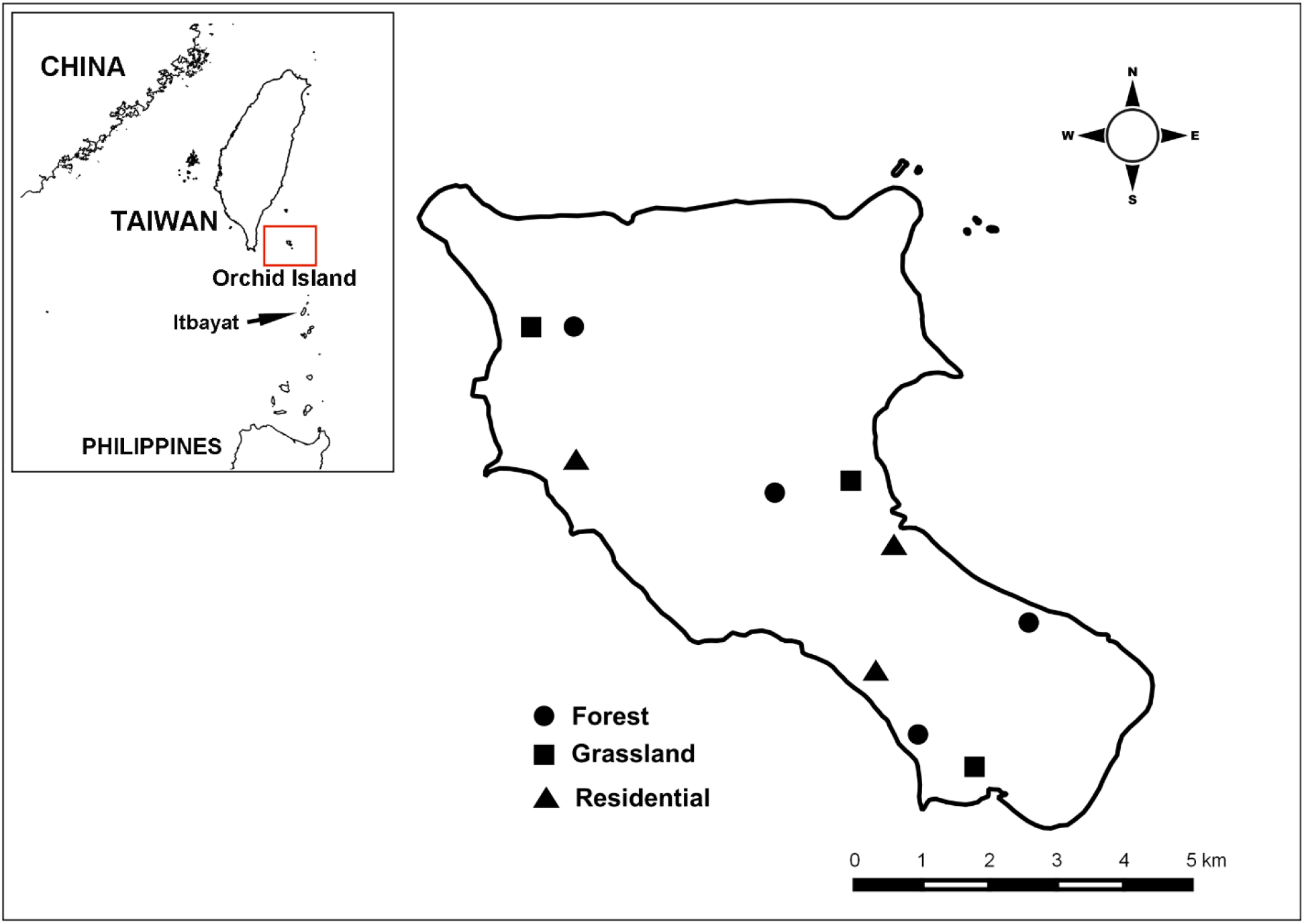
Study sites on Orchid Island of Taiwan.

### 2.2. Small mammal trapping and collection of chiggers

From September 2017 to June 2018, small mammal traps were set up in each of the 10 study sites every three months. In each site, Sherman traps (26.5 × 10.0 × 8.5 cm) and mesh traps (27 x 16 x 13 cm) were deployed alternatively every 10-15 meters along a transect line for a total of 20 traps (each with 10 traps). Preliminary results revealed a much higher capture success of the Asian house rat by using mesh than Sherman traps. However, difficulty in transporting the bulky and unfoldable mesh traps had limited the number of their usage. Foldable Sherman traps were therefore added to increase the trapping effort. This will also allow a comparison of trapping efficacy between the two trap equipment, a useful information for rat eradication program when needed. Traps were baited with sweet potatoes smeared with peanut butter and open for three consecutive nights. For each seasonal trapping session, 10 sites were surveyed within 10 days to avoid any confounding temporal influence on rodents and chiggers.

Once trapped, small mammals were transferred to clean nylon bags and species was identified. Because all trapped rodents belonged to the exotic Asian house rat, they were not released back to the field. An overdose of 0.1 ml of Zoletil 50 anesthetic (Virbac. Carros, France) was injected subcutaneously, followed by cervical dislocation to euthanize sedated rats. Sex and reproductive status of each rodent were determined, and lengths of body, tail, ear, and hind foot (mm) along with body mass (g) were measured. We checked the presence of ectoparasites by thoroughly examining whole body of the animal with the naked eye. Ears with chiggers attached were detached and placed in vials. Chiggers were allowed one day to release themselves from the rat before being preserved in 75% ethanol. We stored chiggers at −20°C for subsequent molecular determination.

### 2.3. Chigger identification

Due to the large number of collected chiggers (>100,000), only one-tenth of chiggers from each rodent was randomly retrieved for species identification. Chiggers were soaked in deionized water 2-3 times to replace ethanol with water and then slide-mounted in Berlese fluid (Asco Laboratories Ltd, Manchester, U.K.). Chiggers were examined under an upright microscope (Olympus BX53, Tokyo, Japan) and identified to species following published keys (Wang and Yu, 1992; Li et al., 1997; Chung et al., 2015).

### 2.4. OT detection in chiggers

Due to the small size of chiggers, a pool of 30 chiggers of the same species from the same rat individual was pooled to retrieve enough DNA for detection of OT. This will also allow potential comparison of infective rate of OT in previous studies that pooled the same number of chiggers (e.g. Kuo et al., 2012; Wei et al., 2020). We followed Kuo et al. (2012) in using the nested polymerase chain reaction (PCR) to target the well conserved 56-kDa type specific antigen located on the OT outer membrane. Laboratory OT strains and nuclease free water were used as positive and negative controls, respectively.

### 2.5. Statistical analysis

Difference in trapping success was assessed with chi-square test. Generalized estimating equations (GEE) was applied to compare number of rodents and their total infested chiggers in each sampling site among habitats and months. Habitat, month, and interactions of both factors were the fixed factors, with site as the subject, and each seasonal sampling as a repeated measures within the site (ten sites, each with four sampling sessions, so overall 40 samples). A normal distribution function and negative binomial log link function were used for rodent and chigger abundance, respectively. Significance of difference was determined based on 95% Wald confidence interval of estimated marginal means.

Generalized linear mixed model (GLMM) was used to analyze how load of chiggers of individual rodent varied with habitat, month, and body size. Habitat, month, interactions of both factors, as well as rodent body weight were the fixed factors, site as a random factor. Similarly, a negative binomial link function were used, and significance of difference was determined based on 95% Wald confidence interval of estimated marginal means.

Mean and 95% confidence interval (CI) of individual-level (per chigger) infection prevalence of OT in chiggers were estimated with a frequentist approach assuming perfect test, with confidence intervals calculated based on binomial theory following Cowling et al. (1999).

All the analyses were performed in SPSS Statistics version 19.0 (IBM Corp.), package lme4 in R 3.6.1 and EpiTools Epidemiological Calculators (Sergeant, 2018).

### 2.6. Ethical statement

All the animal handling procedures were approved by the National Taiwan Normal University Animal Care and Use Advisory Committee (permit number NTNU-106025; NTNU-106048), which adheres to Guideline for the Care and Use of Laboratory Animals established by the Council of Agriculture, Taiwan.

## 3. Results

### 3.1. Rodent abundance across habitats and months

A total of 254 rodents were captured out of 2,400 trap-nights, with a trapping success of 10.6% (number of rodents per trap-night). Only nine out of the 254 rodents were trapped by Sherman traps (trapping success 0.8%) while the remaining 245 rodents were trapped by mesh traps (20.4%). There was a significant difference in trapping success between the two trap types (χ^2^ = 243.2, df = 1, *p* < 0.001). All rodents were the Asian house rat (*R. tanezumi*).

Rodent abundance varied among habitats (χ^2^ = 20.5, df = 2, *p* < 0.001) and months (χ^2^ = 22.2, df = 3, *p* < 0.001), and there was an interaction between habitats and months (χ^2^ = 50.6, df = 6, *p* < 0.001). There were significantly more rodents in forests than in residential sites for most of the months (except for June) (*p* < 0.05). By contrast, rodent abundance was largely similar between forests and grasslands (except for March), and between grasslands and residential sites (except for September) (*p* > 0.05) (Fig. 3A).

**Fig. 3.**
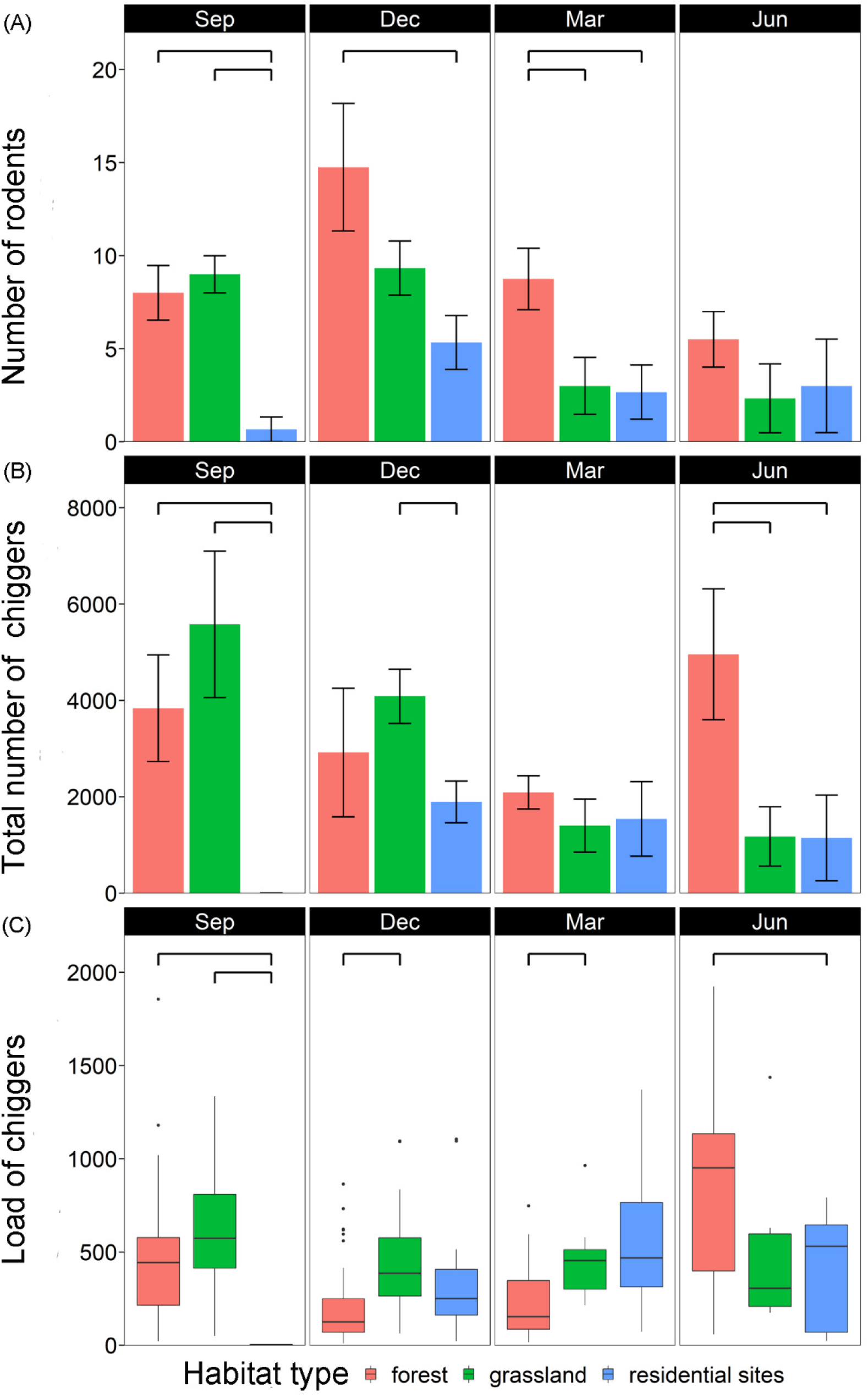
Monthly variations among habitats in (A) number of rodents per study site; (B) number of total chiggers per study site; (C) load of chiggers per rodent individual. Each bar is represented by mean ± SE; significant difference (*p* < 0.05) between two groups is annotated with a bridge.

### 3.2. Total chiggers across habitats, months, and association with rodent abundance

A total of 105,680 chiggers were collected, and all rodents were found infested with chiggers (prevalence = 100%). However, total number of chiggers per site varied among habitats (χ^2^ = 126.5, df = 2, *p* < 0.001) and months (χ^2^ = 50.4, df = 3, *p* < 0.001), and there was an interaction between habitats and months (χ^2^ = 3775.3, df = 6, *p* < 0.001). There were significantly more chiggers in forests than in residential sites in September and June (both *p* < 0.05). There were also more chiggers in grasslands than in residential sites in September and December (both *p* < 0.05). Abundance of chiggers was similar in forests and grasslands except for June (*p* > 0.05) (Fig. 3B).

To further explore whether difference in chiggers was the direct effect of habitats and months, or was indirectly mediated through their effect on rodent abundance (as in *3.1*), rodent abundance was included in the above model as a covariant. Results showed that in addition to the direct effect of habitats, months, and their interactions on total chiggers (χ^2^ = 70.9, 114.0, 2458.4, respectively, all *p* < 0.001), rodent abundance was also positively associated with chigger abundance (χ^2^ = 51.6, β = 0.18, *p* < 0.001) (Fig. 4A). Therefore, habitats and months not only directly but also indirectly affected chigger abundance mediated through their effect on rodent abundance.

**Fig. 4.**
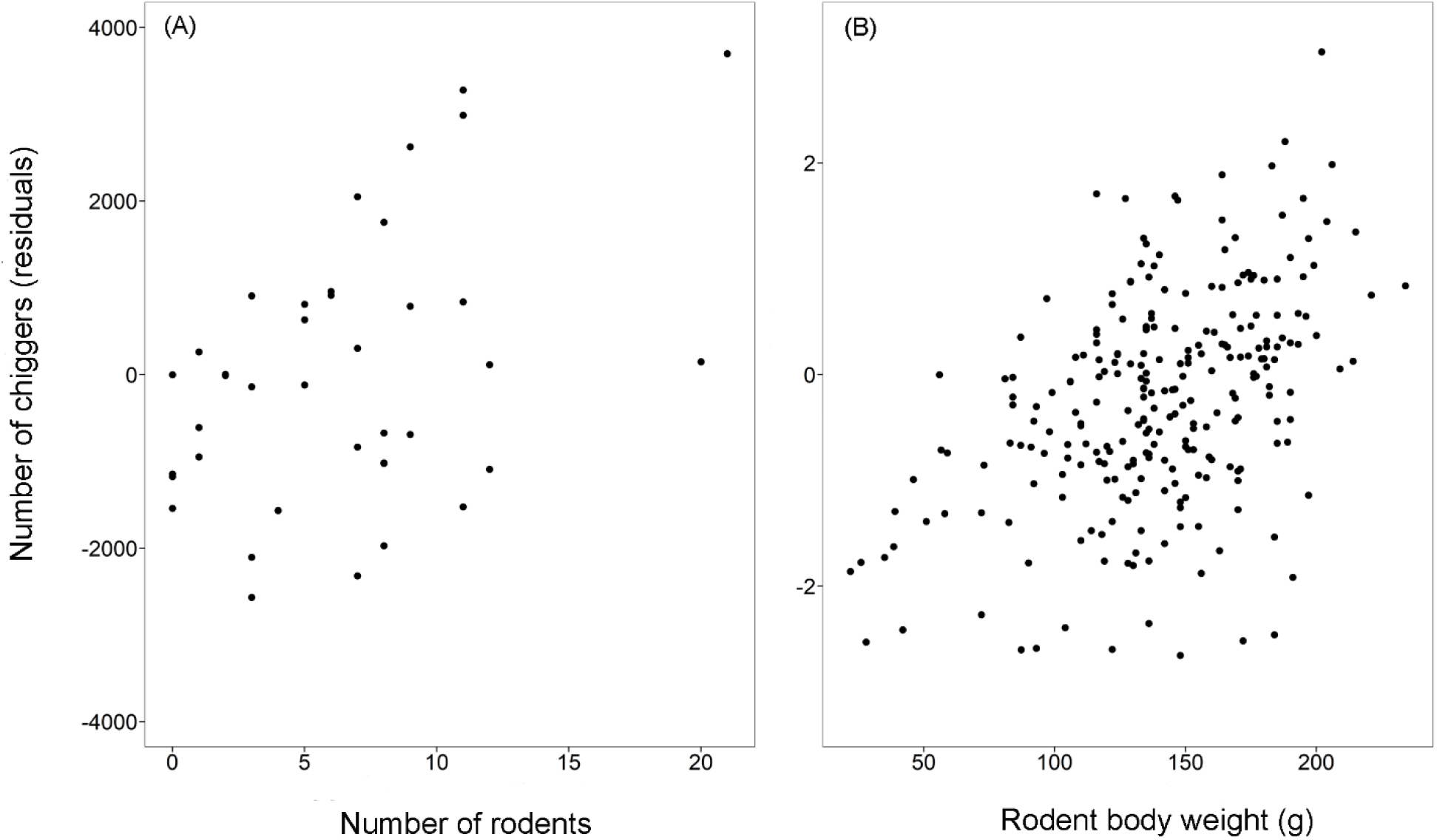
Scatterplots of number of chiggers (residuals, controlled for effect of habitat, month, and their interactions) against (A) number of rodents per study site; (B) rodent body weight. Both scatterplots show significantly positive associations (see the Results).

### 3.3. Chigger load across habitats, months, and association with rodent body weight

Load of chiggers of individual rodent varied among habitats (χ^2^ = 11.9, df = 2, *p* < 0.005) and months (χ^2^ = 80.6, df = 3, *p* < 0.001), and there was an interaction between habitats and months (χ^2^ = 83.2, df = 6, *p* < 0.001). Chigger load was also positively associated with rodent body weight (χ^2^ = 88.8, df = 1, *p* < 0.001) (Fig. 4B). Rodents had higher chigger load in grasslands than in forests in December and March (both *p* < 0.05); chigger load was also higher in grasslands in residential sites in September (*p* < 0.05). Additionally, chigger load was higher in forests than in residential sites in September and June (both *p* < 0.05) (Fig. 3C).

### 3.4. Species composition of chiggers

We morphologically identified 10,368 chiggers to species level. Excluding 208 chiggers that cannot be reliably identified (due to damaged body parts or inadequate placement of specimens), the remaining 10,160 chiggers included 9712 *Leptotrombidium deliense* (95.6%), 341 *Walchia xishaensis* (3.4%), and 107 *Eutrombicula wichmanni* (1.1%)*. L. deliense* was observed in all habitat types across all sampling months. *W. xishaensis* mainly occurred in forests in September and June while *E. wichmanni* was found primarily in a residential site in northwest Orchid Island in March and June.

### 3.5. Prevalence of OT infection in chiggers among chigger species, habitats, and months

A total of 200 pools of chiggers were assayed for OT infection, including 22 pools of *W. xishaensis* and 178 pools of *L. deliense*. OT was not detected in any of the *W. xishaensis* pools (0% prevalence), while prevalence of individual-level (per chigger) infection in *L. deliense* was 1.53% (1.18–1.95%, 178 pools, 37.1%—representing 95% CI, number of pools tested, and pool-level prevalence).

Infection prevalence of *L. deliense* was similar in residential (0.88%, 0.35–1.82%, 30 pools, 23.3%—representing mean, 95% CI of prevalence, number of pools tested, and pool-level prevalence), grassland (1.53%, 0.96–2.30%, 62 pools, 37.1%) and forest sites (1.79%, 1.24–2.48%, 86 pools, 41.9%). Likewise, there was no seasonal difference in prevalence among September (1.55%, 0.86–2.57%, 40 pools, 37.5%), December (2.08%, 1.37–3.00%, 62 pools, 46.8%), March (1.45%, 0.82–2.36%, 45 pools, 35.6%), and June (0.71%, 0.26–1.55%, 31 pools, 19.4%).

## 4. Discussion

In comparison to residential sites, forests harbored more rodents and chiggers. In addition, infection prevalence of OT in *L. deliense* chiggers, the most important scrub typhus vector species in Southeast Asia (Kawamura et al., 1995) and also the primary vector species in Orchid Island, in forests was two times that of residential sites (1.79% vs. 0.88%). We also found a positive association between rodent body weight and chigger load, as well as between rodent abundance and total chiggers.

In Orchid Island, the expansion of commensal Asian house rat from residential sites to forests and grasslands has helped the spread of chiggers and thus scrub typhus infection risks from human settlements to these two natural habitats. This is particularly prominent in the forests. Forests had more rodents than the other two habitats; moreover, larger differences occurred during the winter and early spring (December and March, Fig. 3A) when the monsoon brought in high wind and low temperature and when food resources were presumably more scarce. The forests, with complex stand structures and abundant tree species that bear large fruits, such as *P. formosanum* and *P. pinnata*, could provide rodents a better shelter.

At the same time, forests also harbored significantly more chiggers (all months pooled, forests (mean chiggers per site: 3282) higher than grassland (2475), in turn than residential sites (273), both *p* < 0.05). More strikingly, we observed a great increase in chigger abundance in June when chiggers in the other habitats were in decline (Fig. 3B). Higher rodent number and intrinsic difference in habitat characteristics have both contributed to more chiggers in the forests (section *3.2*). Higher rodent abundance should increase the probability of host finding of questing chiggers that are themselves vulnerable to desiccation (Traub and Wisseman, 1974; Muul et al., 1977; Kawamura et al., 1995) when forests, at the same time, can maintain a higher soil humidity with their closed canopies. Moreover, tropical forest soils are rich in arthropods (Sayer et al., 2010; Cole et al., 2016) which provide nymphal and adult mites necessary food to feed on. Our study not only validates potential occurrence of scrub typhus in the primary forests (Traub and Wisseman, 1974) but also further demonstrates forests as a better habitat for chiggers.

On the other hand, habitat difference in chigger load is more varied (Fig. 3C). In most months, chigger load were not always lower in residential sites than the other two habitats. In March, for example, mean chigger load was much higher (although not significantly higher) in residential sites than in forests. This can be due to much more rodents in the latter habitat (Fig. 3A) so that chiggers therein were less concentrated among rodent hosts; total chigger abundance was still higher in the forest (Fig. 3B). Unpaved areas surrounding the human settlements might help sustain the chigger population. Importantly, the occurrence of large number of chiggers in residential sites warrants caution of scrub typhus infection.

As previously reported (Perkins et al., 2003; Kiffner et al., 2011; Mysterud et al., 2015), we also observed a positive association between body size and loads of ectoparasites, so that larger rodents generally carried more chiggers (Fig. 4B). However, it should be stressed that this study included only island but not mainland populations. Whether chigger load will increase with subsequent gigantism when rodents colonize islands can therefore not be validated here. It has been corroborated that skull size of *R. tanezumi* in Orchid Island is significantly larger than that in mainland Taiwan (Jenq, 2007). The next step is to verify whether the increased body size is accompanied with higher chigger load following the colonization.

Likewise, we found more chiggers in those sites with more rodents (Fig. 4A). Again, this does not necessary mean that such positive association will hold when rodent population increases from mainland to island. To our knowledge, no estimation of population density or trapping success of *R. tanezumi* in mainland Taiwan has ever been reported. Any valid comparison will further be confounded by the type of traps used, as evidenced in this study that using Sherman instead of mesh traps will detect a much lower rodent population. The reality that chigger abundance can increase with rodent density, nevertheless, indicates that a comparison of rodent density and chigger abundance between mainland and island is a promising and worthwhile effort.

It has been observed that exotic animals are less frequently parasitized and also parasitized with fewer species in introduced than in their native ranges (Torchin et al., 2003). A few mechanisms have been proposed to explain such phenomenon. Because distribution of parasites among hosts are highly aggregated, with most individuals lightly parasitized (the so called 20/80 rule) (Woolhouse et al., 1997), most native parasites will be left behind in introduced animals. For native parasites with complex life cycle, introduced regions might lack hosts necessary for finishing their life cycle; or parasites of introduced regions might be host specific or haven’t yet evolved to utilize newly introduced hosts (Torchin et al., 2003). However, *R. tanezumi* trapped in this study were all infested with chiggers (prevalence = 100%); some individuals were even parasitized with close to 2,000 chiggers. Because host is required in only a single life stage (larva) of trombiculid mite, and the *L. deliense* chigger can utilize a very wide range of host species (Stekolnikov, 2021), introduced hosts can be infested with no fewer chiggers, particularly when rodents were introduced from habitats unfavorable for chiggers (paved areas on mainland) to their favorable habitats (forest on island). Unfortunately, chigger infestation on *R. tanezumi* in mainland Taiwan has never been reported, especially areas surrounding seaports. A surveillance of ectoparasite infestation on *R. norvegicus*, also a commensal rodent, in Kaohsiung seaport has found no chiggers, suggesting few if any chigger in the seaport (Yan, 2011). The enemy release hypothesis posits that the success of invaded species can be partially attributed to their being less parasitized in invaded sites compared to the original ranges (Keane and Crawley, 2002; Mitchell and Power, 2003; Torchin et al., 2003). This may not hold, nevertheless, when the parasite is of simple life cycle, a host generalist, and is transferred from unsuitable to suitable habitats.

It is unknown whether chiggers in Orchid Island are introduced or are native to Orchid Island. Rodents are the primary hosts of chiggers (Harrison and Audy, 1951; Traub and Wisseman, 1974; Kawamura et al., 1995), including *L. deliense* (Stekolnikov, 2021). If chiggers are introduced, whether accompanied *R. tanezumi* or other exotic host species, their abundance will increase following raised number and habitat expansion of *R. tanezumi*. Similarly, when chiggers are native, their number will not greatly increase until after the introduction of rodents, which are a more suitable host.

Likewise, the origin of *R. tanezumi* in Orchid Island has not been fully resolved. The island is inhabited mainly by aboriginals of Polynesian origin known as the Tao tribe (also called Yami). Tao is suggested descendants of indigenous residents from the Itbayat Island of the Philippines, 145 km south of Orchid Island (Fig. 2), because both speak the same language (de Beauclair, 1959), although recent molecular data showed few genetic exchange between the two tribes (Loo et al., 2011). Therefore, even though Orchid Island is closer to Taiwan, *R. tanezumi* might instead be introduced from the Itbayat Island; this species also occurs in the Batanes Islands that include Itbayat Island (Gonzalez et al., 2008). Genetic data, however, showed that *R. tanezumi* in Orchid Island is more closely related to Taiwan mainland than to the Philippines despite that the Orchid Island population is not recently descended from Taiwan mainland (Jenq, 2007). Therefore, current data still support closer origin from Taiwan mainland.

Our study stresses the importance of further investigation on vector-borne and zoonotic diseases on islands. For example, some common commensal species, such as *R. tanezumi* and *Rattus rattus*, can not only adapt to diverse environments (Chaisiri et al., 2015; Shiels et al., 2017; Wilson et al., 2017) but also host a variety of zoonotic diseases (Chaisiri et al., 2015; Kosoy et al., 2015). They are also regular invaders on islands (Johnson and Stattersfield, 1990). Accordingly, disease risks posed by their introductions to island inhabitants should be carefully evaluated.

## Acknowledgements

This study was financially supported by Taiwan Ministry of Science and Technology (MOST 104-2314-B-003-002-MY3; MOST 105-2621-M-003-003) awarded to CCK. The funders had no role in study design, data collection and analysis, decision to publish, or preparation of the manuscript

